# A chromosome-scale genome assembly of the flax rust fungus reveals the two unusually large effector proteins, AvrM3 and AvrN

**DOI:** 10.1101/2025.04.28.651126

**Authors:** Jana Sperschneider, Jian Chen, Claire Anderson, Emmanuelle Morin, Xiaoxiao Zhang, David Lewis, Eva Henningsen, Igor V. Grigoriev, John P. Rathjen, David A. Jones, Sebastien Duplessis, Peter N. Dodds

## Abstract

Rust fungi comprise thousands of species many of which cause disease on important crop plants. The flax rust fungus *Melampsora lini* has been a model species for the genetic dissection of plant immunity since the 1940s, however the highly fragmented and incomplete reference genome has so far hindered progress in effector gene discovery. Here, we generate a fully-phased, chromosome-scale assembly of the two nuclear genomes of *M. lini* strain CH5, resolving an additional 320 Mbp of sequence compared to the previous short-read assembly. The 482 Mbp dikaryotic genome is at least 79% repetitive with a large proportion (∼40%) of the genome comprised of young, highly similar transposable elements. The assembly resolves the known effector gene loci some of which carry complex duplications that were collapsed in the previous assembly. Using a genetic map followed by manual correction of gene models, we identify the *AvrM3* and *AvrN* genes which encode unusually large fungal effector proteins and trigger defense responses when co-expressed with the corresponding resistance genes in *Nicotiana tabacum*. We locate the genes linked to the tetrapolar mating system on chromosomes 4 and 9, but in contrast to the cereal rusts which have one pheromone receptor gene per haplotype, in flax rust three pheromone receptor genes are found with two of them closely linked on one haplotype. Taken together, we show that a high-quality assembly is crucial for resolving complex gene loci and given the increasing number of fungal effectors of large size, the commonly applied criterion for effector candidates as being small proteins as needs to be reconsidered.

## Introduction

The rust fungi encompass over 7,000 species which together form the single largest group of plant pathogens (Aime and McTaggart 2021). They are responsible for diseases on some of the most important agricultural crops (wheat, oat, maize, barley, coffee, foxtail millet and soybean) as well as on timber species and numerous native plants of ecological significance. Rust fungi are obligate biotrophs with complex life cycles. Some rusts complete their life cycle on one host (autoecious) whereas others require two alternating unrelated hosts (heteroecious). The genus *Melampsora* contains both autoecious and heteroecious species and most heteroecious *Melampsora* rusts complete the telial stage on poplar or willow leaves. In contrast, the flax rust fungus *Melampsora lini* is autoecious and completes its life cycle on one host plant, namely flax (*Linum usitatissimum*). The autoecious life cycle of *M. lini*, ease of flax rust disease scoring and favorable flax plant growth habits that allow sequential infection as well as the capacity for genetic transformation (Lawrence et al. 2010) meant that *M. lini* became a model for the study of rust fungi.

The flax rust system has been used as a model for the genetic dissection of plant disease resistance since the 1940s, providing the basis of the gene-for-gene relationship concept (Flor 1942; Dodds 2023). In this model, absence or inactivity of either a plant resistance (*R*) gene or the corresponding pathogen avirulence (*Avr*) effector gene leads to disease establishment. The first *Avr* gene identified in a rust fungus was *AvrL567*. This gene was cloned using a cDNA marker cosegregating with the avirulence phenotype in an F2 *M. lini* family, followed by examination of a 26.5-kbp genomic DNA region from the avirulence locus (Dodds et al. 2004). Based on the observation that *AvrL567* is expressed in haustoria and delivered into the plant cell, a targeted search for genes encoding small, secreted proteins that were expressed in a haustorium-specific cDNA library led to the identification of *AvrM, AvrP, AvrP123* and *AvrP4* (Catanzariti et al. 2006; Barrett et al. 2009). Taken together, prior to the availability of an *M. lini* genome sequence, four *Avr* loci (*AvrL567, AvrM, AvrP/AvrP123, AvrP4*) encoding proteins ranging from 92 to 377 amino acids (aas) in size were identified.

With the availability of high-quality genome assemblies, effector identification has moved from reference-free approaches (such as map-based cloning) to *in silico* prediction of effector candidates. This approach has two limitations: first, it relies heavily on the quality of the genome assembly and its effector gene annotations; and second, it can produce long lists of effector candidates of up to thousands of genes that require validation. Thus, effector candidate selection routinely includes prioritization through population genomics analysis (e.g. genome-wide selection analysis, genome-wide association studies/quantitative trait locus mapping) or by genomic compartmentalization analysis (e.g. accessory chromosomes, repeat-rich regions). An alternative to candidate prioritisation is high-throughput screening, which has recently become possible through a major technical advance that enables small pools of candidate genes (Wilson et al. 2024) or large libraries of hundreds of candidates (Arndell et al. 2024; Chen et al. 2025) to be screened by expression in plant protoplasts. Again, a highly accurate genome assembly and effector gene annotation are pre-requisites as candidate effector libraries are synthesized.

Whilst population genomic analysis has been successfully applied to fungal pathogens with haploid genomes and high genetic diversity (Kanja and Hammond□Kosack 2020; Plissonneau et al. 2017), this is precluded in most rust fungi due to the presence of long-lived clonal populations. One notable exception is *M. lini* for which a high-density genetic map is available which can be used for effector gene identification (Lawrence et al. 1981; Anderson et al. 2016). However, we thus far lack a high-quality genome assembly for *M. lini*. Whilst high-quality, chromosome-scale assemblies for fungal pathogens with small, haploid genomes are now easily attainable, rust genomes have been more difficult to resolve due to their relatively large size, high repeat content and dikaryotic life stages during which they carry two haploid nuclear-separated genomes that are challenging to phase. The current 162.8 Mbp short-read genome assembly of the *M. lini* CH5 isolate, an F1 hybrid of *M. lini* strains C and H (Lawrence et al. 1981), is highly fragmented in 21,130 scaffolds with 14% gaps (189.5 Mbp genome assembly size including gaps; 16,271 gene models; 46% repetitive sequences) (Nemri et al. 2014). It also represents a mixed, collapsed version of the two haploid genomes from the two distinct C and H parental nuclei. The high-density genetic map for CH5 resulted in 27 linkage groups (Anderson et al. 2016) but only 69% of the short-read genome assembly could be anchored onto the map and 190 incorrectly assembled, chimeric scaffolds were identified based on substantial differences between genetic and physical maps. Despite the poor genome assembly, *AvrL2* and *AvrM14* were isolated using markers from the genetic map combined with amplification of alleles missing from the assembly using strain H and strain C genomic DNA (Anderson et al. 2016). However, although the genetic map contained markers that co-segregated with *AvrM3* and *AvrN*, no gene candidates could be identified (Anderson et al. 2016) due to limitations of the assembly. Here we present an improved assembly for *M. lini* strain CH5 based on PacBio HiFi and Hi-C sequencing, which resolves the two C and H sub-genomes to chromosome-level. We show that together with *in planta* expression data for gene model correction, this allows for identification of *AvrM3* and *AvrN*.

## Results

### A chromosome-scale, nuclear-assigned genome assembly of *M. lini* resolves an additional 320 Mbp of sequence

We generated and assembled PacBio HiFi and Hi-C reads from *M. lini* strain CH5 and arrived at 36 chromosomes with 18 centromeres visible for each haplotype in the Hi-C contact maps (Table 1, Supplementary Figure 1, Figure 1A). The 18 haplotype C chromosomes have a total size of 233.3 Mbp with only two gaps and the 18 haplotype H chromosomes have a total size of 237.5 Mbp with only seven gaps. The 18 chromosomes in each haplotype show high BUSCO completeness of 96%. All unplaced contigs are less than 347 kbp in size (239 unplaced contigs of total size 11.5 Mbp, N/L50: 67/50.4 kbp) and are highly enriched for repetitive sequences especially ribosomal RNA repeats (Table 1). To independently assess the nuclear phasing of the C and H haplotypes generated from Hi-C data, we extracted the sequences of the C and H alleles of 13,412 RADseq markers mapped in the CH5F2 family (Anderson et al. 2016) (Supplementary Table 1) and aligned these to the assembled chromosomes, recording only perfect alignments. For the C-allele marker sequences, 13,410 (99.99%) have a perfect hit to the C haplotype and only 123 (0.92%) also have a perfect hit to the H haplotype. For the H-allele marker sequences, 13,412 (100%) have a perfect hit to the H haplotype and only 83 (0.62%) also have a perfect hit to the C haplotype. Thus, the genetic data for this genome-wide set of markers confirms that the assembly is fully-phased and correctly nuclear-assigned (Table 1), independently verifying the accuracy of Hi-C data for nuclear assignment in dikaryons.

**Table 1:** *M. lini* genome assembly statistics for the C and H haplotypes as well as for the unplaced contigs.

|  | C haplotype | H haplotype | Unplaced contigs |
| --- | --- | --- | --- |
| Number of scaffolds | 18 | 18 | 239 |
| Assembly size (Mbp) | 233.31 | 237.46 | 11.49 |
| GC content | 40.3% | 40.3% | 38.9% |
| Complete BUSCOs | 96% | 95.8% | 0.2% |
| Duplicated BUSCOs | 0.7% | 0.6% | 0.1% |
| Number of genes | 15,556 | 16,630 | 159 |
| Repetitive sequence | 77.99% | 78.36% | 95.27% |
| % retroelements | 63.83% | 63.87% | 13.31% |
| % DNA transposons | 3.88% | 3.97% | 1.23% |
| % rRNA | 0.07% | 0.09% | 68.45% |
| Aligned bases between haplotypes | 98.09% | - |  |
| Average identity of 1-to-1 alignments | 99.26% | - |  |
| Average identity of M-to-M alignments | 98.93% | - |  |
| Total SNPs between haplotypes | 931,075 | - |  |
| C markers with perfect alignment | 99.99% | 0.92% | 0.1% |
| H markers with perfect alignment | 0.62% | 100% | 0.9% |
| Number of telomeres | 16 forward, 15 reverse | 17 forward, 14 reverse | 1 forward |

**Figure 1:**
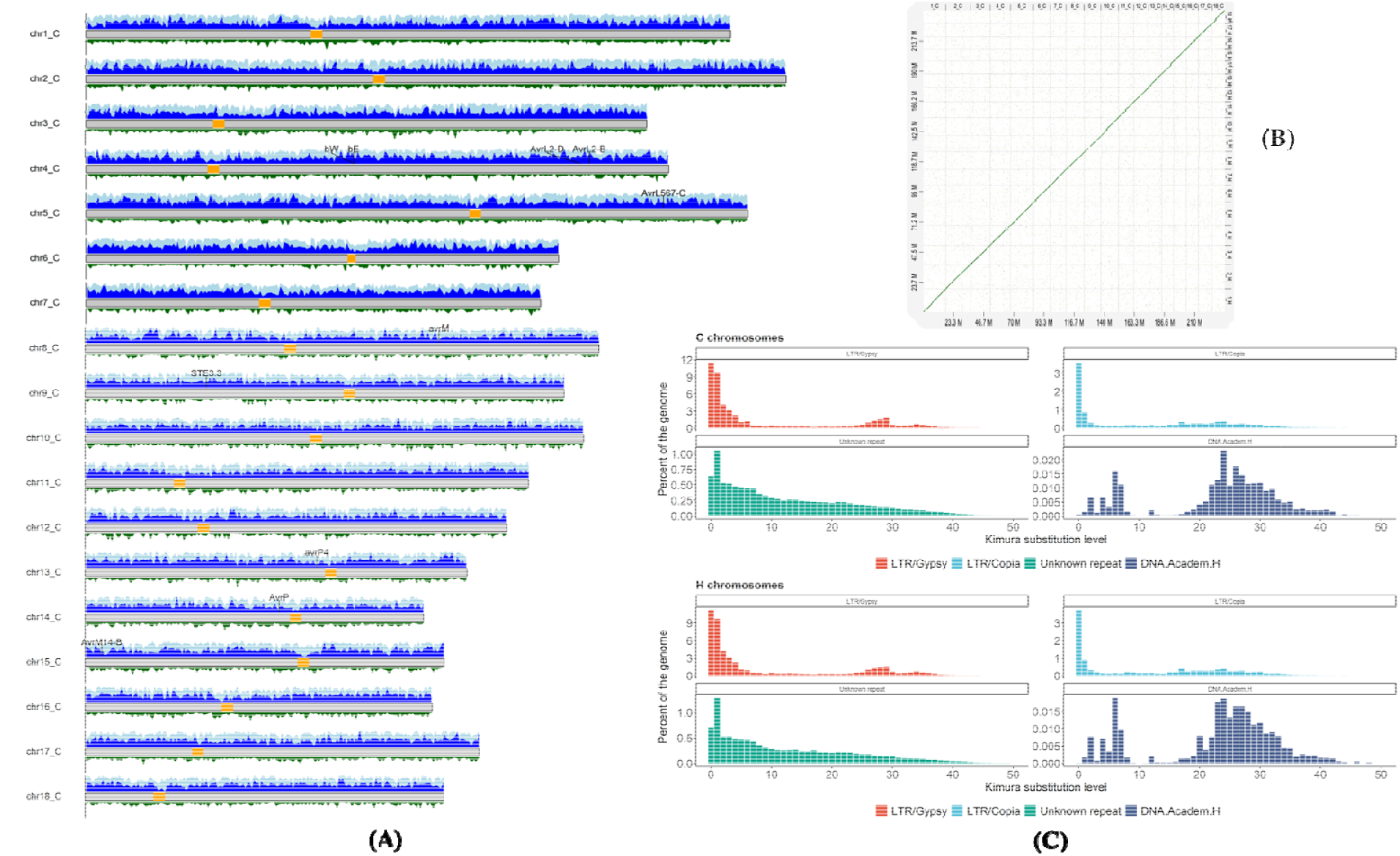
The *M. lini* haplotype chromosomes. **(A)** The C haplotype chromosomes are shown with the location of the centromeres (yellow regions), effector genes and mating-type genes on chromosomes 4 and 9. Repeat elements, young repeat elements and annotated genes are shown as coverage tracks in light blue, dark blue and green, respectively (50 kbp windows). **(B)** Synteny plot between the C (x-axis) and H (y-axis) chromosomes generated in D-Genies (Cabanettes and Klopp 2018). **(C)** Kimura substitution levels for four repeat classes are shown. Young TEs copies that are highly conserved have < 5% Kimura substitution levels.

The haplotype C and H chromosomes share ∼99% sequence identity (Table 1) with only one large structural rearrangement, an inversion of ∼2 Mbp on chromosome 4 and a region with no alignment on chromosome 9, which corresponds to the pheromone receptor mating-type locus (Figure 1B and see below). The C and H chromosomes carry 15,556 and 16,630 annotated genes, respectively, and are at least 78% repetitive with a total of 368 Mbp of repeat sequence. Over 64% of the genome is covered by retroelements with ∼ 45% belonging to the LTR Gypsy/DIRS1 class. We observed that 40.2% of the genome is comprised of repetitive elements with low sequence divergence (< 5%), indicative of a recent genome expansion in *M. lini* through young repeat elements (Figure 1C). The centromeres are less repetitive (67%) than the remainder of the genome, with only 22% of the centromeric repeats constituting young repeat elements. Taken together, the *M. lini* genome is a vast improvement over the previous short-read assembly and resolves the chromosomal landscape and centromeres of the two haplotypes with an additional 320 Mbp of sequence and 15,915 additional genes.

### Most of the known *Avr* loci of *M. lini* are complex duplications but some are highly conserved

Thus far *Avr* genes from six loci have been cloned from *M. lini* strain CH5, some of which occur in complex genomic regions that were either collapsed, chimeric or both in the previous short-read assembly (Dodds et al. 2004; Catanzariti et al. 2006; Barrett et al. 2009; Nemri et al. 2014; Anderson et al. 2016). All of these *Avr* loci are now fully resolved in the assembly. The most complex genomic region is the *AvrM* locus which had been described to contain at least 5 paralogs separated by not less than 10 kbp (*AvrM-A* to *-E*) for the avirulence allele on the H haplotype and a single gene, *avrM*, in the virulence allele on the C haplotype (Catanzariti et al. 2006). The previous short-read assembly collapsed all of these gene variants into a single gene (Nemri et al. 2014) whereas this assembly resolves eight AvrM paralogs, seven on the H haplotype and one on the C haplotype (Figure 2A). However, the gene annotation pipeline only annotated the virulence allele *avrM* on the C haplotype, but none of the seven *AvrM* paralogs on the H haplotype. We used RNA-seq evidence to manually annotate the seven *AvrM* genes on the H haplotype. These encode the previously described AvrM-A (1 copy), AvrM-B (1 copy), AvrM-C (three copies) and AvrM-D (two copies) gene variants. However, the AvrM-E variant described by Catanzariti et al. (2006) was not present. This sequence was detected in a single EST clone that contained a mutation leading to a premature stop codon and it is possible that this was an artefact introduced during cDNA cloning. In the assembly, the *AvrM* paralog regions share a common, highly repetitive composition defined by the presence of a repeated sequence element containing a DNA/hAT-Ac transposon, a gene encoding a hypothetical protein (983 aas; only annotated on the C haplotype), an AvrM gene, a LINE/TAD1 repeat element and an ORF related to a UbiA prenyltransferase (UbiA) (not annotated and covered by annotated repeat family) (Figure 2). The right-most copy of the AvrM repeat on the H haplotype carries a UbiA pseudogene (frameshift). Taken together, the assembly resolves the complex H haplotype *AvrM* locus that contains 7 paralogs spanning ∼285 kbp.

**Figure 2:**
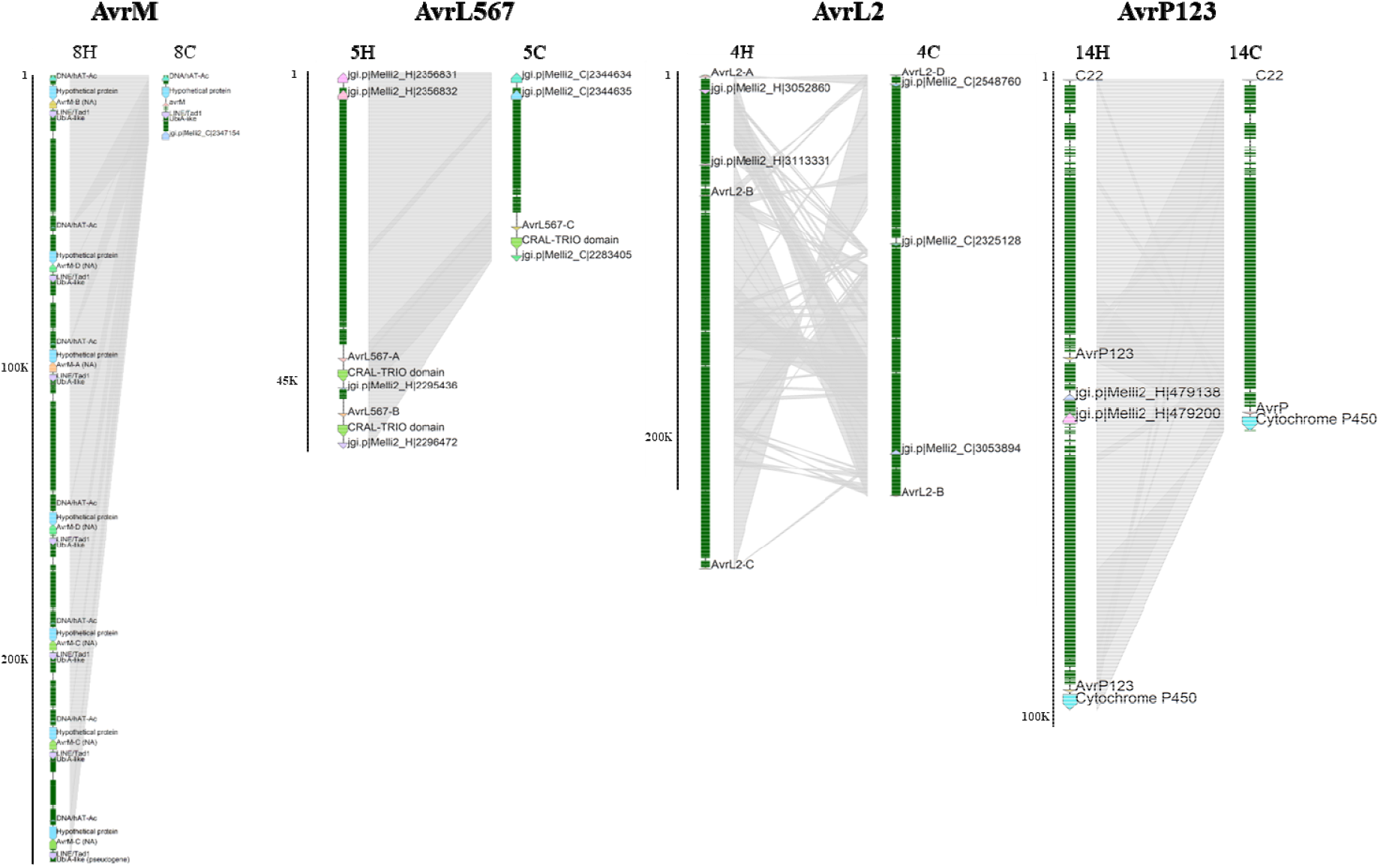
Structural variation at *Avr* loci in CH5. Figure shows synteny plots between the C and H haplotypes at the *AvrM, AvrL567, AvrL2* and *AvrP123* loci. Genes are labelled with their identifier and NA stands for ‘not annotated’ in the gene annotation. Repetitive elements are shown in green boxes. Separate scale bars are shown to the left of each locus. Shaded boxes connect haplotype regions of conserved sequence.

The *AvrL567* locus on the H haplotype contains the duplicated avirulence effector genes *AvrL567-A* and *AvrL567-B*, as expected. The virulence allele *AvrL567-C* is found on the C haplotype. As described by Dodds *et al*., 2004, the AvrL567-A and -B genes on the H haplotype are in a duplicated 26.5 kbp region that is surrounded by LTR Gypsy elements and a region with a high frequency of recombination that is also found in the C haplotype of this locus. We also note a highly repetitive LTR Gypsy element-rich region preceding the C allele in the assembly. *AvrL2* was previously identified through map-based cloning, but the *AvrL2* locus was not resolved in the short-read assembly due to its complexity (Nemri et al. 2014) and PCR amplification was used to confirm the sequences of the *AvrL2* variants in each haplotype (avirulent: *AvrL2-A, AvrL2-B, AvrL2-C*; virulent: *AvrL2-B, AvrL2-D*) (Anderson et al. 2016). Our assembly places *AvrL2* within the large inversion of ∼2 Mbp on chromosome 4, consistent with the very low recombination rate around this locus that was reported previously (Anderson et al. 2016). The C haplotype and H haplotype *AvrL2* loci span ∼77 kbp and ∼248 kbp, respectively, with complex structural rearrangements yet correct assembly of the five *AvrL2* genes (Figure 2C). Rust strain CH5 contains two alleles at the *AvrP123* locus with the *AvrP* allele conferring avirulence to the *P* resistance gene in flax and the *AvrP123* allele conferring recognition by *P1, P2* and *P3* (Barrett et al. 2009). The assembly correctly placed *AvrP* on the C haplotype on chromosome 14, next to a putative cytochrome P450 monooxygenase gene, while there is a previously undetected duplication containing two identical copies of the *AvrP123* gene in the H haplotype (Figure 2C). A previously described pseudogene (C22) of related sequence (Barrett et al. 2009) occurred on both haplotypes adjacent to *AvrP*/*AvrP123*.

In contrast to the complex *AvrM, AvrL567, AvrL2* and *AvrP123* loci, the *AvrP4* and *AvrM14* loci are highly conserved regions with no gene duplication (Supplementary Figure 2). The *AvrM14-A* avirulence and *AvrM14-B* virulence alleles are correctly annotated on the H and C haplotype, respectively, whilst they were collapsed into one sequence in the previous assembly (Nemri et al. 2014). These genes are in a highly conserved region close to multiple LTR Gypsy and Copia elements (Supplementary Figure 2). Similarly, the *AvrP4* locus is highly conserved between the two haplotypes (Supplementary Figure 2). Taken together, the *Avr* loci are all fully resolved in their expected copy numbers and were all correctly annotated, apart from the seven *AvrM* genes on the H haplotype and the two copies of *AvrL2-B* where the N-terminus is missing six amino acids resulting in a shorter signal peptide sequence.

### Genetic map enables the identification of the effector proteins AvrM3 and AvrN

Co-segregating genetic markers have been reported for the effector genes *AvrM3* and *AvrN*, but the revious short-read assembly did not allow accurate candidate gene identification (Anderson et al. 2016). We mapped the two reported markers co-segregating with *AvrM3* and two flanking markers onto the chromosomes, which identified a 164 kbp target region on chromosome 9C and a 201 kbp target region on chromosome 9H. On the H haplotype, which carries the avirulence allele (Anderson et al. 2016), we identified one candidate secreted protein of 715 aas, with the gene model supported by *in planta* RNA-seq reads and the protein predicted as a cytoplasmic effector by EffectorP 3.0 (Sperschneider and Dodds 2022). However, the corresponding allele on the C haplotype was not annotated correctly according to an inspection of the RNA-seq read alignments. We manually re-annotated the C haplotype gene which also encodes a 715 aa secreted protein (Supplementary Figure 3) with 12 aa differences, all of which are located in the C-terminal region (Figure 3A). The locus structure is highly conserved between the two haplotypes (Supplementary Figure 4) and was collapsed into one contig in the previous assembly (Nemri et al. 2014).

**Figure 3:**
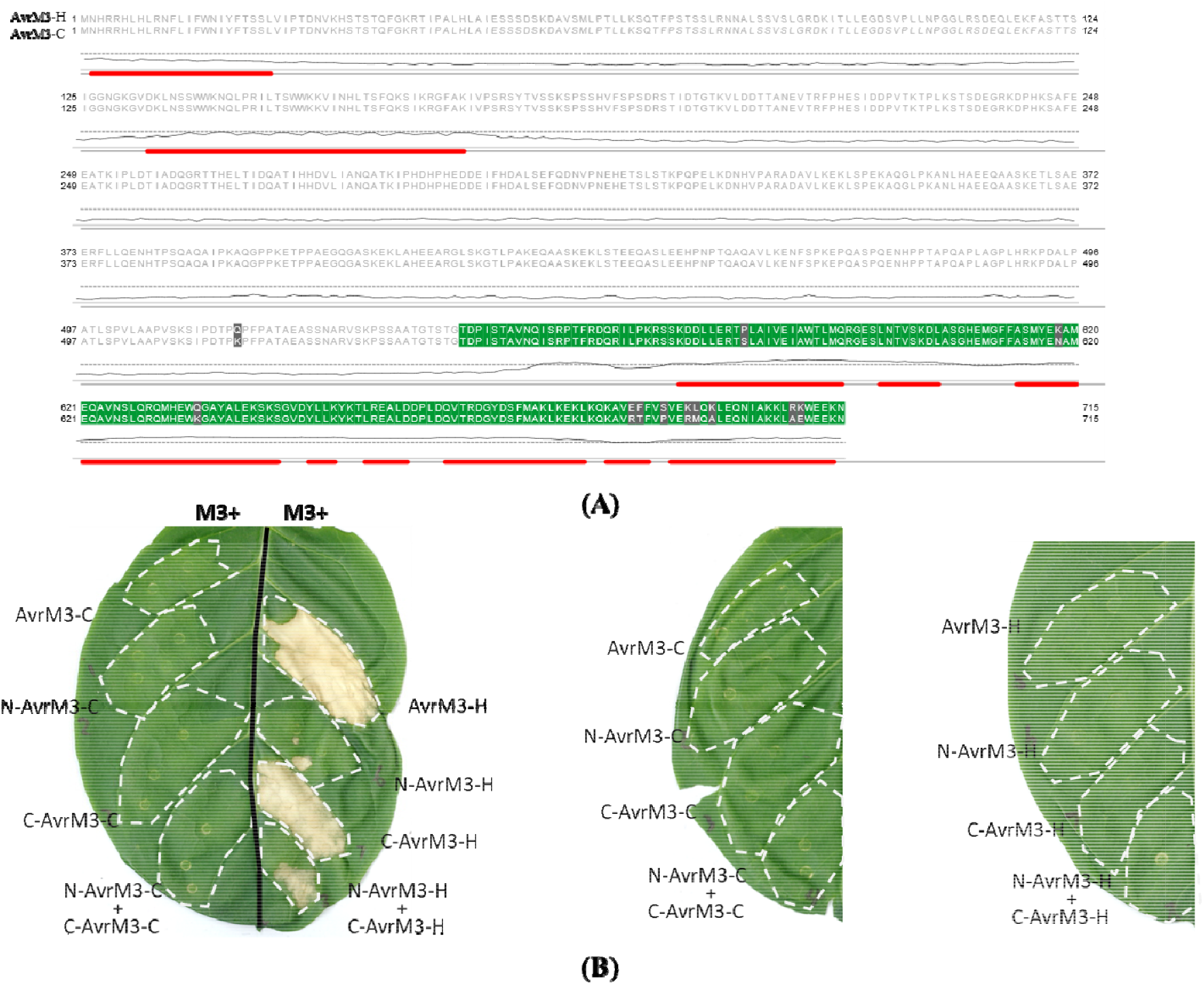
AvrM3 sequence alignment and agroinfiltration assays. **(A)** Sequence alignment of AvrM3-C and AvrM3-H. Alpha helices in the AlphaFold3 predicted structure are highlighted as red boxes and pLDDT values of prediction confidence are shown as a graph, with pLDDT of 70 indicated as a dotted line. Sequence differences between the avirulence and virulence alleles are highlighted with gray boxes and the C-terminal truncation region is shown in green. **(B)** Transient coexpression in *N. tabacum* of M3 with 3HA tag at the C-terminus with AvrM3-C and AvrM3-H and their truncated versions fused with a YFP tag at the C terminus (left leaf). Middle and right leaves show expression of the AvrM3-C and H proteins and fragments in the absence of M3.

Agroinfiltration assays in *Nicotiana tabacum* showed cell death induction upon transient co-expression of M3 with the AvrM3-H variant but not AvrM3-C, confirming specific recognition (Figure 3B). An AlphaFold3 model of AvrM3-H confidently predicted alpha-helical structures (pLDDT score > 70) in the C-terminal region, but the remainder of the protein was unstructured with low confidence (Figure 3A). We generated genes encoding N-terminal and C-terminal fragments of AvrM3-H and AvrM3-C split at amino acid 544 which is N-proximal to the predicted alpha-helices (Figure 3A). Agroinfiltration assays showed that co-expression of M3 with the N-terminal fragment of AvrM3-H induced no response, whereas the C-terminal fragment of AvrM3-H with M3 induced strong cell death. However, none of the AvrM3-C fragments triggered cell death with M3 in *N. tabacum*. This suggests that the C-terminal structured region of AvrM-H is sufficient for recognition, with the amino acid differences in this region determining the difference in recognition of AvrM3-H and -C proteins.

For *AvrN*, we mapped the six reported co-segregating markers and two flanking markers onto the CH5 genome, which identified a 232 kbp target region on chromosome 8C and a 241 kbp target region on chromosome 8H. The region on chromosome 8C contained a single candidate gene encoding a secreted protein of 559 aas that is predicted as a cytoplasmic effector by EffectorP 3.0 (Sperschneider and Dodds 2022) and the gene model is supported by alignments of RNA-seq reads from infection. However, visual inspection of the annotation and alignments of RNA-seq reads led to the manual re-annotation of the allelic gene on the H haplotype, which contains a pre-mature stop codon leading to a truncated protein of 191 aas due to an LTR Copia repeat insertion (Figure 4A, Supplementary Figure 4). This fits the observation that the C haplotype carries the avirulence allele and the H haplotype the virulence allele (Anderson et al. 2016). Agroinfiltration assays in *N. tabacum* showed cell death induction upon transient co-expression of the avirulence effector AvrN-C with the N protein, but not of the truncated virulence allele AvrN-H (Figure 4B). The avirulence effector AvrN-C co-immunoprecipitated with the N resistance protein whereas the virulence effector AvrN-H and the negative control AvrSr35 did not show interaction with N (Figure 4C). This confirms that this secreted protein encoded by the C haplotype is indeed AvrN. Again, the two effector genes *AvrN-C* and *AvrN-H* were collapsed into one contig in the previous assembly (Nemri et al. 2014).

**Figure 4:**
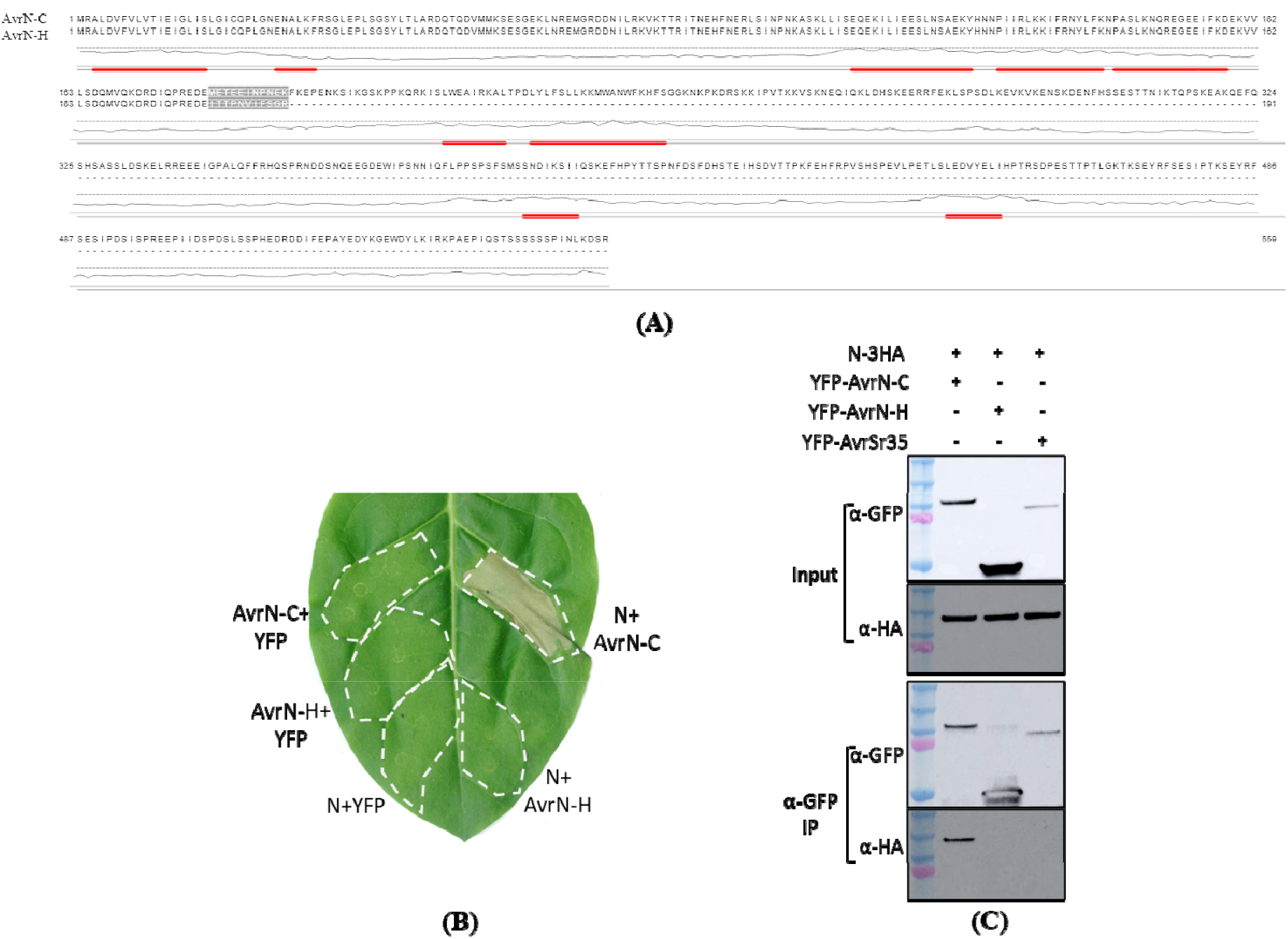
AvrN sequence alignment and agroinfiltration assays. **(A)** Sequence alignment of AvrN-C and AvrN-H. Alpha helices in the AlphaFold3 predicted structure are highlighted as red boxes and pLDDT values of prediction confidence are shown as a graph, with pLDDT of 70 indicated as a dotted line. Sequence differences between the avirulence and virulence alleles are highlighted with gray boxes followed by a truncation in the virulence allele. **(B)** Transient coexpression of N-3HA with AvrN-C and AvrN-H fused to a YFP tag at the N-terminus in *N. tabacum*. Leaves were documented 3 days after infiltration. **(C)** Co-IP assays of N and the AvrN effectors. AvrN-C and AvrN-H fused with YFP tag were used to pull down N-3HA. AvrSr35 was used as a negative control.

### AvrM3 and AvrN are unusually large fungal effector proteins

Both the AvrM3 and AvrN proteins share sequence homology with proteins only from other *Melampsora* species. We found homologs with ∼70% amino acid identity encoded in the genome of *M. larici-populina* (Duplessis et al. 2011) and with ∼50% amino acid identity in the genome of *M. americana* (Crowell et al. 2022). The *AvrN* gene has one intron whilst the *AvrM3* gene has an unusually high number of 17 introns (Supplementary Figures 3 and 5). Furthermore, the AvrN and AvrM3 proteins are unusually large for fungal effectors at 559 and 715 aas, respectively. We plotted the length distribution of 165 published fungal effector proteins and over 75% of these are < 300 aas (Supplementary Table S2, Figure 5A). Only eleven fungal effectors, including AvrN and AvrM3, are outliers with protein length > 500 aas (Figure 5A). Lastly, we assessed if the *M. lini* effector genes share commonalities in expression in germinated spores and in flax leaves collected at 2, 3, 4, 5, 6, 7 and 8 dpi. Most effector genes show a similar expression profile with lowest expression in germinated spores and highest expression at 2-5 dpi (Figure 5B). However, *AvrM3* shows a different expression pattern with relatively high expression in early infection but also in germinated spores. No significant differences in expression were found between the avirulence and virulence alleles for *AvrM3* or any of the other effector genes (data not shown).

**Figure 5:**
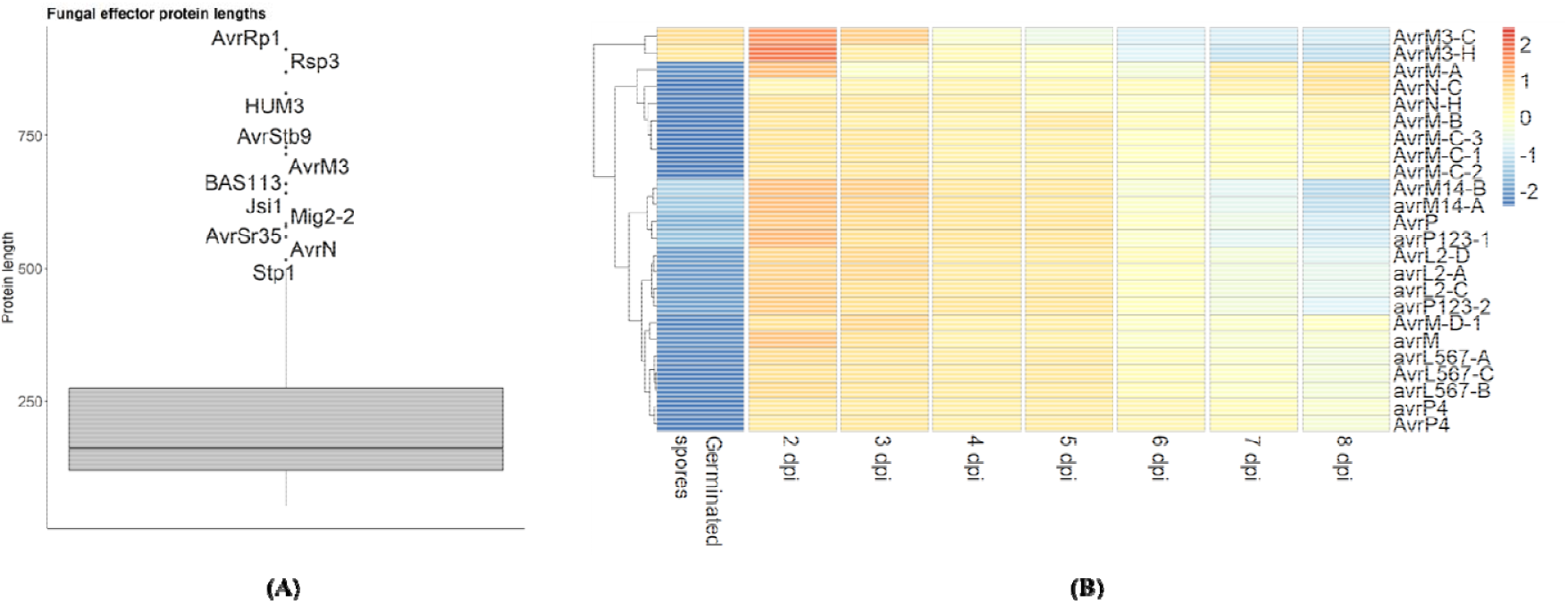
Fungal effector protein size and expression of effector genes. **(A)** Distribution of the protein lengths of 165 published fungal effectors is shown. Only eleven effectors, including AvrN and AvrM3, are outliers with protein lengths of over 500 aas. **(B)** Heatmap showing expression levels (log(TPM) expression values scaled by row) for *M. lini* effector genes in germinated spores and infected leaves from 2 to 8 days post infection (dpi).

### The *M. lini* chromosomes carry recombination cold spots linked to the mating-type loci

We used the genetic map (Anderson et al. 2016) to study the distribution of recombination on the chromosomes (Figure 6A). Recombination rate analysis highlighted high variation along the chromosomes, with an average recombination rate of 26.1 cM/Mbp (Figure 6B). We observed recombination cold spots at sub-telomeric regions as well as low recombination rates at the centromeres and in regions of structural variation (Figure 6C). For example, chromosome 4 features a large inverted region of 2 Mbp that is a recombination cold spot and contains *AvrL2*. However, on average the genomic regions containing the known *Avr* genes do not show significant differences in recombination rate to all other genes (Figure 6C). Genomic regions with low recombination rates are significantly more repetitive than those with high recombination rates (Figure 6D).

**Figure 6:**
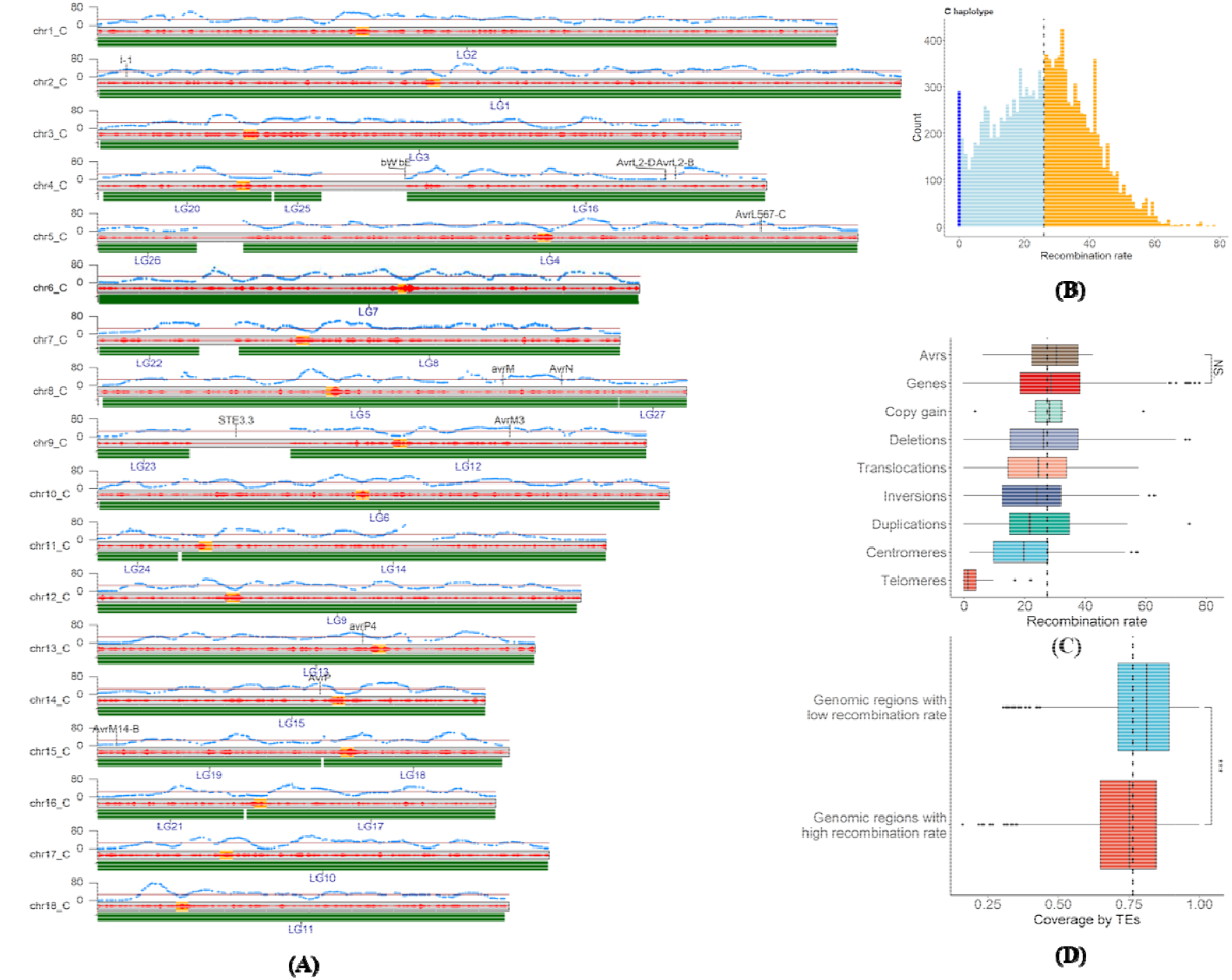
Recombination rate analysis for the C chromosomes. **(A)** The C chromosomes are shown with the location of the linkage groups (LGs) defined by Anderson *et al*. (2016) shown by green bars, centromere locations shaded in yellow, graphs showing the density of genetic markers in 50 kbp windows (red) and the recombination rate (blue line). The mean recombination rate is shown as a red line. **(B)** The frequency (*y*-axis) of recombination rates (*x*-axis) on the C chromosomes is shown. Recombination cold spots are shown in dark blue, low recombination rates are shown in light blue and high recombination rates are shown in orange, with the mean recombination rate indicated by a dotted line. **(C)** Distribution of recombination rates in intervals constituting centromeres, sub-telomeric regions (50 kbp at start and end of chromosome), structural variants, genes and *Avr* effector genes. (**D**) Distribution of repeat element (TE) coverage (*x*-axis) in regions with low or high recombination rates as defined in (B).

Ten of the 27 linkage groups in the genetic map (Anderson et al. 2016) span complete chromosomes (Figure 6A). A further four chromosomes are completely covered by pairs of linkage groups separated by small gaps. However, four chromosomes (4, 5, 7 and 9) each contain a large region (0.94-2.4 Mbp) with no genetic markers (Figure 6A). We speculated that these regions might contain the mating type loci because we expect strong suppression or absence of recombination in these regions, which would prevent the detection of segregating co-dominant genetic markers that were the basis of the genetic map. In rust fungi the two known mating-type loci are the pheromone receptor (PR) locus, which contains pheromone receptor (*STE3*) genes and pheromone precursor (*mfa*) genes, and the homeodomain (HD) locus, which contains two divergently transcribed homeodomain transcription factor genes called *bE* (*b*East) and *bW* (*b*West) (Cuomo et al. 2017). Indeed, two of the four marker-free regions on chromosomes 4 and 9 (2.04 Mbp and 2.4 Mbp, respectively) contain the previously characterized mating-type genes. We inspected the gene annotations of the other two marker-free regions on chromosomes 5 and 7 (1.1 Mbp and 0.94 Mbp, respectively) but did not find a link to previously characterized mating-type genes.

We first inspected the PR locus on chromosome 9. We located four pheromone receptor genes *STE3*.*2*.*1-STE3*.*2*.*4* using the homologs from *M. larici-populina* which were located on three different scaffolds in that assembly (Duplessis et al. 2011). We note that we had to correct the gene model for *STE3*.*2*.*3* using the RNA-seq data. The *STE3*.*2*.*1* genes are located on chromosomes 1C and 1H and have one amino acid difference between the two haplotypes. The *STE3*.*2*.*3* gene is located on chromosome 9H (3.75 Mbp) and the *STE3*.*2*.*2* and *STE3*.*2*.*4* genes are located 58 kbp apart in a repetitive region on chromosome 9C at 3.28 and 3.22 Mbp, respectively (Figure 7A). A phylogenetic tree placed both *STE3*.*2*.*2* and *STE3*.*2*.*4* in a clade with *STE3*.*2*.*2* from Puccinia species and separate from *STE3*.*2*.*3* or *STE3*.*2*.*1* (Figure 7B), suggesting a duplication of this gene in the *Melampsora* lineage. The *STE3*.*2*.*3* and *STE3*.*2*.*4* genes display very similar expression patterns, with low expression in germinated spores and increasing expression during the infection time course, with a peak of expression at 8 dpi. In contrast, the *STE3*.*2*.*2* gene which is closely linked to *STE3*.*2*.*4* has lower expression levels during infection, but higher expression in germinated spores possibly suggesting a diverged function (Figure 7C). We detected only negligible expression levels for the *STE3*.*2*.*1* genes.

**Figure 7:**
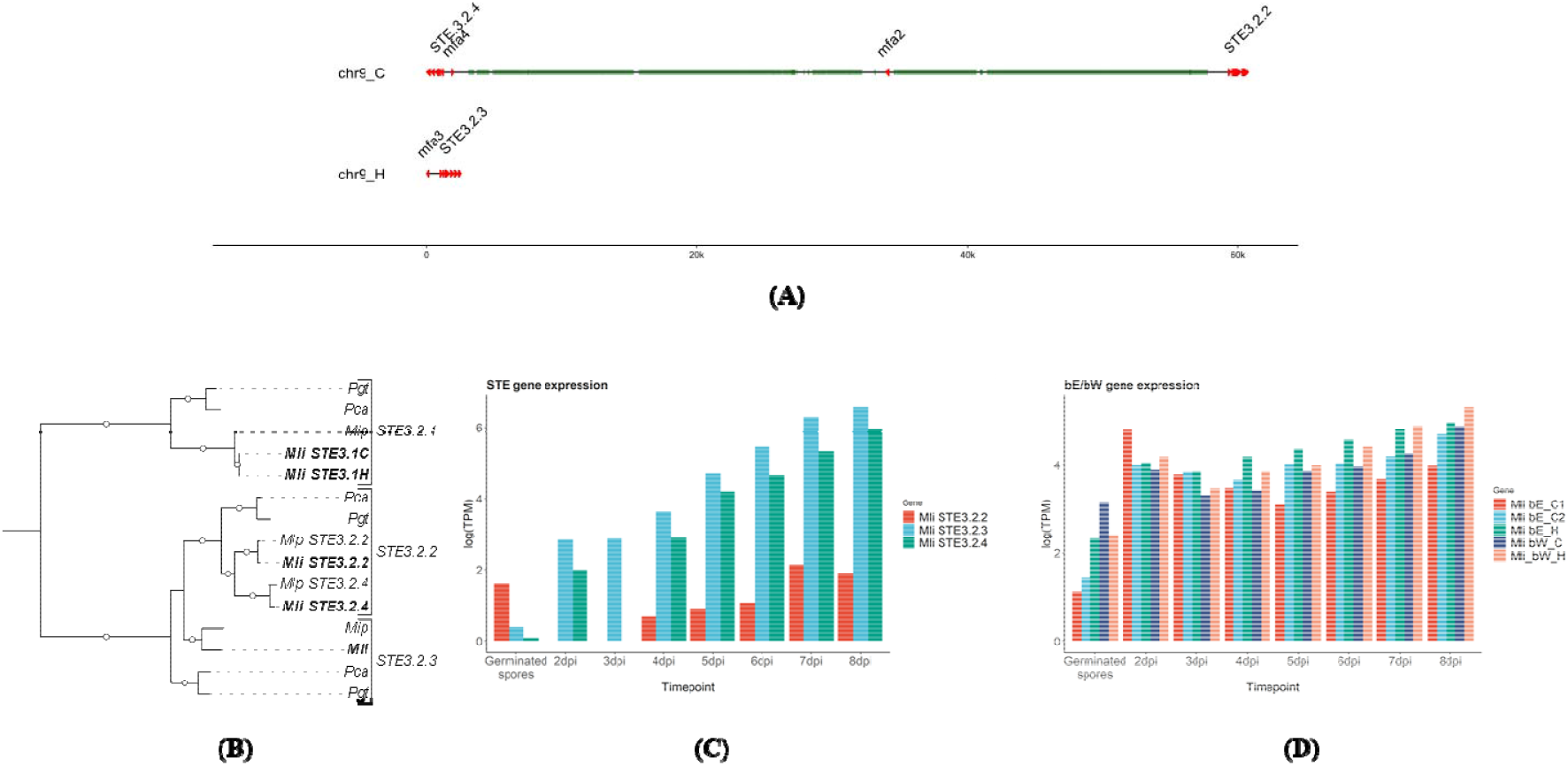
Analysis of the mating-type locus in flax rust. **(A):** A duplication of a STE3 allele on haplotype C is shown. Repetitive elements are shown in green. **(B):** A phylogeny of the STE3 proteins shows the duplication of the STE3.3.2 clade alleles in both *M. lini* and *Melampsora larici-populina*, whereas the cereal rusts *Puccinia graminis* f. sp. *tritici* (*Pgt*) and *Puccinia coronata* f. sp. *avenae* (*Pca*) only carry one STE3.2.2 allele and one STE3.2.3 allele. **(C)** Expression of the *STE* genes and **(D)** expression of the *bE/bW* genes.

We then located the closest *mfa* genes to the *STE3*.*2* genes. On chromosome 9C, the *STE3*.*2*.*4* gene is closely linked to an *mfa4* gene encoding a 39 aa protein that is only 601 bps away (Figure 7A). On the same chromosome, the *STE3*.*2*.*2* gene is 25 kbp away from an *mfa2* gene that encodes a 79 aa protein. On chromosome 9H, the *STE3*.*2*.*3* gene is 895 bps away from an *mfa3* gene that encodes a 54 aa protein. Inspection of RNA-seq alignments confirmed that *mfa4* and *mfa3* are both highly expressed during infection, whereas *mfa2* shows lower expression (data not shown). Taken together, the *STE3*.*2*.*4* gene on haplotype C and the *STE3*.*2*.*3* gene on haplotype H share close resemblance in expression and proximity to an *mfa* gene whereas the *STE3*.*2*.*2* gene on haplotype H has lower expression and is more distant to a lowly expressed *mfa* gene that is located in a highly repetitive region.

Lastly, we inspected the HD locus on chromosome 4, which contains two linked HD transcription factor genes (*bE*-*HD1* and *bW*-*HD*2) (Cuomo et al. 2017). We first corrected the gene model for *bE* on haplotype C (termed *bE-C*) with the RNA-seq data. This also showed the presence of two isoforms for that gene (*bE*-C1 and *bE*-C2) that differ by one amino acid with similar expression levels (Figure 7D). The bE proteins in *M. lini* are of similar length (C haplotype: 398/399 aas and H haplotype: 392 aas) to those in the cereal rusts (*Puccinia graminis* f. sp. *tritici* and *Puccinia coronata* f. sp. *avenae*: 373 aas). In contrast, the bW proteins in *M. lini* are shorter (C haplotype: 456 aas and H haplotype: 458 aas) than those in the cereal rusts (*Puccinia graminis* f. sp. *tritici*: 618 aas and *Puccinia coronata* f. sp. *avenae*: 572 aas). However, all of the bE and bW proteins contain the expected DNA binding motif sequences (WFXNXR) (Kües et al. 2011).

## Discussion

A telomere-to-telomere genome assembly is the foundation for comprehensive studies of the biological function of genes in an organism and its genome biology. Furthermore, in non-haploid organisms fully-phased assemblies are critical for accurate comparisons between haplotypes and to assess their genetic variation. Here, we report the first fully-phased, telomere-to-telemere assembly for a rust fungus of the *Melampsora* genus, the CH5 isolate of *M. lini*. Early estimations of nuclear DNA content by flow cytometry estimated that the nuclear DNA content of *M. lini* was 2.5 times that of *Puccinia graminis* f. sp. *tritici* (Eilam et al. 1994). The haploid genome size for *P. graminis* f. sp. *tritici*. is ∼85 Mbp (Li et al. 2019) which would imply a 213 Mbp haploid size for *M. lini*. The first genome assembly for *M. lini* isolate CH5 used Illumina sequencing data (read lengths: ∼300-5000 bps), leading to an assembly size of 189.5 Mbp, only 89% of the expected haploid genome size (Nemri et al. 2014). The high fragmentation of the assembly (14.1% N’s in the assembly, longest scaffold size: 239.7 kbp, L50: 31.5 kbp) indicated that the genome carries a high number of repetitive sequences that the short sequences could not accurately assemble (Nemri et al. 2014). Using PacBio HiFi and Hi-C sequencing data, we show that the haploid genome size for *M. lini* is ∼235 Mbp, which is very large for a fungal pathogen, but agrees well with the estimate from relative nuclear fluorescence. Analysis of the HiFi assembly indicates the reasons for the poor Illumina assembly. First, the two haplotypes are highly similar at 99% sequence identity, which would lead to collapsed assembled regions if the short Illumina reads do not span sequence differences. Second, we observed a large proportion of young repetitive elements of the LTR class that are relatively long in sequence, which again would lead to collapsed regions using the short Illumina reads. This recent genome expansion in *M. lini* through young repeat elements was recently dated to have occurred in multiple recent bursts over a few million years in contrast to continuous repeat accumulation in e.g. *Phakopsora pachyrhizi* or *P. graminis* f. sp. *tritici* (Corre et al. 2025)

The *M. lini* HiFi genome is a vast improvement over the previous short-read assembly and resolves the chromosomal landscape of the two haplotypes, an additional 320 Mbp of sequence and 15,915 additional genes. These improvements are particularly impactful for the accurate assembly of complex gene loci such as those that encode effectors. For example, we show that the *AvrM* effector genes on the H haplotype occur in a cluster of 7 paralogs spanning 285 kbp which is larger than the longest assembled scaffold in the previous assembly (Nemri et al. 2014). Furthermore, the C and H haplotype *AvrL2* loci span ∼77 kbp and ∼248 kbp, respectively, which is larger than the L50 of the previous assembly (50.4 kpb). However, we note that despite the inclusion of high-quality, stranded RNA-seq data at eight time points several *Avr* gene models as well as some of the highly conserved mating-type genes had to be manually annotated or corrected. This underlines that comparative genomics studies should include steps to search for genes that are present in the genome yet missed or mis-annotated by gene annotation pipelines.

Using the *M. lini* genetic map (Anderson et al. 2016) followed by manual assessment and correction of gene models resulted in the identification of *AvrM3* and *AvrN*. These are two effector genes for which co-segregating markers were reported, yet the previous assembly did not allow effector gene identification in the marker intervals. Here, we locate and validate the two effectors *AvrM3* and *AvrN*, both of which encode proteins of unusually large size for fungal effectors. Fungal effectors are commonly identified by selecting secreted proteins of small size (e.g. < 300 aas) but the increasing identification of larger effectors warrants reconsideration of the practice of stringently applying small size cutoffs. We showed that about 25% of confirmed fungal effectors are over 300 aas and ∼7% are over 500 aas. Given that many fungal effectors have been identified in screens that excluded larger proteins, the predominance of smaller effectors may be affected by some ascertainment bias, with larger effector under-represented.

Rust fungi have been suggested to carry a tetrapolar mating system where two physically unlinked genomic regions, the P/R or *a* locus and the HD or *b* locus, control non-self-recognition (Cuomo et al. 2017). We confirmed that unlike the cereal rusts which have only one copy of the *STE3*.*2* gene, there appears to be a duplication of this gene in *M. lini* with both genes (*STE3*.*2*.*4* and *STE3*.*2*.*2*) located in close proximity. Recombination suppression on chromosomes 4 and 9 extends well beyond the mating-type loci as a 2 Mbp region in line with some other fungi (Hartmann et al. 2021). Experimental data from selfing and intercrossing of the two *M. lini* strains CH5 and I indicated that it may possess a more complicated system than a bi-allelic (+/) *a* locus and a multi-allelic *b* locus (Lawrence 1980). Additional genomic data from other *M. lini* strains is necessary to test this hypothesis.

## Materials and Methods

### PacBio HiFi DNA and Hi-C sequencing

High molecular DNA from urediniospores was extracted as previously described (Schwessinger and Rathjen 2017; Li et al. 2019). DNA quality was assessed with a Nanodrop spectrophotometer (Thermo Scientific) and the concentration quantified using a broad-range assay in a Qubit 3.0 fluorometer (Invitrogen). DNA library preparation (10–15□kbp fragments Pippin Prep) and sequencing in PacBio Sequel II Platform (One SMRT Cell 8M) were performed by the Australian Genome Research Facility (AGRF) (St Lucia, Queensland, Australia) following manufacturer guidelines. For DNA crosslinking and subsequent Hi-C sequencing, 100□mg of urediniospores was suspended in 4□ml 1% [v/v]formaldehyde, incubated at room temperature for 20□min with periodic vortexing. Glycine was added to 1.0□g per 100□ml and the suspension was centrifuged at 1,000□*g* for 1□min and the supernatant was removed. Spores were then washed with H_2_O, centrifuged at 1,000□*g* for 1□min and the supernatant removed. The spores were then transferred to a liquid nitrogen-cooled mortar and ground before being stored at −80□°C or on dry ice. After treatment, spores were shipped to Phase Genomics (Seattle, Washington, USA) for Hi-C library preparation and sequencing (restriction enzymes: DpnII, DdeI, HinfI, MseI).

### Genome assembly and chromosome curation

The HiFi reads were assembled using hifiasm 0.16.1 in Hi-C integration mode and with default parameters (24.5□Gb HiFi reads and 14□Gb Hi-C reads) (Cheng et al. 2022). Contaminants were identified using sequence similarity searches (BLAST 2.11.0 -db nt -evalue 1e-5 -perc_identity 75) (Altschul et al. 1990). HiFi reads were aligned to the assembly with minimap2 v2.22 (-ax map-hifi –secondary=no) (Li 2018) and contig coverage was called using bbmap’s pileup.sh tool on the minimap2 alignment file (http://sourceforge.net/projects/bbmap/). All contaminant contigs, contigs with less than 5x coverage and the mitochondrial contigs were removed from the assembly. BUSCO completeness was assessed with v.3.0.2 (-l basidiomycota_odb9 -sp coprinus) (Simão et al. 2015). Chromosomes were curated using visual inspection of Hi-C contact maps produced using Hi-C-Pro 3.1.0 (MAPQ□=□10) (Servant et al. 2015) and Hicexplorer 3.7.2 (Ramírez et al. 2018). Chromosomes were numbered with reference to the *Puccinia graminis* f. sp. *tritici* nomenclature (Li et al. 2019) informed by location of BUSCOs. To assess phasing, C and H marker sequences (Anderson et al. 2016) were mapped to the genome allowing no mis-matches using bowtie 1.3.1 (-v0) (Langmead et al. 2009). The two haplotypes were compared to each other with mummer 4.0.0rc1, using nucmer and dnadiff (Marçais et al. 2018).

### Gene annotation, repeat annotation and secretome prediction

Both haplotypes were annotated using the JGI Annotation Pipeline (Grigoriev et al. 2014) which detects and masks repeats and transposable elements, predicts genes using a combination of ab initio, homology-based, and transcriptome-based gene prediction tools, chooses a best gene model at each locus to provide a filtered genes catalog, clusters the filtered catalogs into draft gene families, automatically assigns functional descriptions (such as GO terms, EC numbers), and creates a JGI genome portal with tools for public access and community-driven curation of the annotation (https://mycocosm.jgi.doe.gov/Melli2_CH5). Proteins were predicted as secreted if the presence of a signal peptide was detected with SignalP, with D-cutoff values set to “sensitive” (version 4.1; option eukaryotic; Petersen et al., 2011), and no transmembrane helix or one, with the helix start before the signal peptide cleavage position, found by TMHMM using default parameters (version 2.0; Melén et al., 2003). De novo repeats were predicted with RepeatModeler 2.0.2a and the option -LTRStruct (Flynn et al. 2020) and RepeatMasker 4.1.2p1 (-s -engine ncbi) (http://www.repeatmasker.org) was run with the RepeatModeler library to obtain statistics about repetitive element content. The scripts calcDivergenceFromAlign.pl and createRepeatLandscape.pl were used to extract Kimura divergence values.

### RNA-seq data generation and analysis

An infection series of flax (*Linum usitatissimum*, variety Hoshangabad) inoculated with *M. lini* isolate CH5 spores was prepared and exposed to dark high-humidity conditions (< 20°C) for 24 hrs to encourage *M. lini* spore germination. The remaining infection period occurred in controlled growth conditions (24°C) under 12 h light/dark cycles without ramping. Leaf samples were collected from 2-8 days post inoculation (dpi). Additionally, samples of germinated *M. lini* uredineospores were collected by incubation of the spores on water for 8 hrs at room temperature. All samples consisted of 3 replicates. Total RNA was prepared from these samples using the Maxwell RSC Plant RNA Kit (Promega) and mRNA further purified for sequencing on the Illumina platform by the Ramaciotti Centre for Genomics (UNSW Sydney, Australia), generating 150 bp paired-end reads. Reads were cleaned with fastp version 0.22.0 (--length_required 20) (Chen et al. 2018).

For gene model correction, we aligned the reads of each sample to the reference genome assembly using STAR 2.7.9 in 2-pass mode (--alignIntronMin 5 --alignIntronMax 3000 --alignMatesGapMax 3000 --outFilterMultimapNmax 100) (Dobin et al. 2013). For gene expression analysis, we used Salmon 1.10.1 (Patro et al. 2017) in genome decoy mode to generate TPM values.

### Avr loci synteny plots and effector analysis

To plot synteny at the effector gene loci, we first ran minimap2 2.28 with settings -DP -k19 -w19 -m200 (Li 2018) and used gggenomes (Hackl et al. 2024) for visualization. We used miniprot (Li 2023) to locate the AvrM3 and AvrN proteins in the *M. larici-populina* (Duplessis et al. 2011) and *M. americana* (Crowell et al. 2022) genome assemblies. We used Jalview 2.11.4.1 to visualize protein alignments (Waterhouse et al. 2009) and the predicted AlphaFold3 structures (Abramson et al. 2024).

### Transient expression in *Nicotiana tabacum*

The coding sequences (without the signal peptide) of *Avr* gene alleles were synthesised with Gateway overhangs, cloned into pDONR207 via a Gateway BP reaction, and subsequently into expression vectors (Supplementary Table S3) via Gateway LR reaction (Invitrogen). Plasmids used in this study are listed in Supplementary Table S4. *Nicotiana tabacum* plants seedlings were transplanted into Scotts-Debco soil mixed with osmocote and were grown in a growth room at 23□°C with a 16-hour light period and used for agroinfiltration at the 3-to 4-week-old stage. *Agrobacterium tumefaciens* cultures were grown at 28°C overnight with shaking at 200 RPM in Luria-Bertani liquid medium with appropriate antibiotic selections. Cells were pelleted by centrifugation and resuspended in infiltration mix (10□mM MES pH□5.6, 10□mM MgCl_2_, 150□μM acetosyringone) to a final concentration of OD_600_ = 1.0 and incubated at room temperature for two□hours before infiltration. For documentation of cell death, leaves were photographed or scanned 3 to 5□days after infiltration.

### Protein extraction and co-immunoprecipitation

Co-immunoprecipitation experiments were performed as described (Chen et al. 2023). Total proteins were extracted in 800 μl protein extraction buffer (25 mM Tris-HCl, pH 7.5, 150 mM NaCl, 1 mM EDTA, 10 mM DTT, 1 mM PMSF, and 0.1% NP-40, protease inhibitor cocktail (11836170001, Roche) and 0.5% polyvinylpolypyrrolidone). After two rounds of centrifugation at 4 °C, 600 μL of the supernatant were transferred to a round bottom 2.0 mL tube and an equal volume of protein extraction buffer with 5 μL equilibrated GFP-Trap Magnetic Agarose beads (ChromoTek) were added. Samples were incubated for 60-90 min at 4°C with gentle rotation on a wheel. Magnetic beads were separated and washed with protein extraction buffer and four times with wash buffer (25 mM Tris-HCl, pH 7.5, 150 mM NaCl, 1 mM EDTA). Input and GFP pull down protein samples were separated by SDS-PAGE and then transferred to nitrocellulose membranes for immunoblot assays using mouse anti-GFP (REF 11814460001, Roche Switzerland) and goat anti-mouse-horseradish peroxidase (170–5047, BioRad, USA), and rat anti-HA-horseradish peroxidase antibodies (Roche, REF 12013819001, Switzerland), respectively.

### Recombination and mating type locus analysis

We used MareyMap (sliding window setting) (Rezvoy et al. 2007) to estimate local recombination rates along the genome from the genetic map (Anderson et al. 2016). We annotated structural variation with syri 1.6.3 (Goel et al. 2019) on an alignment between the two haplotypes (minimap2 -ax asm5 –eqx) (Li 2018). We calculated the recombination rate for each feature class using bedtools closest (Quinlan and Hall 2010). We identified centromeres through visual inspection of the Hi-C contact maps. For the mating type genes, we used miniprot (Li 2023) to locate the positions of the *M. larici-populina* homologs on the chromosomes. Gene models at those locations were manually inspected with the RNA-seq alignments. For the phylogenetic tree calculation, we used ClustalW (Larkin et al. 2007) and RAxML v8.2.11 with model PROTGAMMAJTT and 100 bootstraps (Stamatakis 2006).

## Data availability

The genome and annotation are available at https://mycocosm.jgi.doe.gov/Melli2_CH5. The PacBio HiFi, Hi-C, RNA-seq reads and corrected gene models are available at the CSIRO Data Access Portal https://data.csiro.au/collection/csiro:35097 and under the NCBI BioProject PRJNA239538.

## Acknowledgements

We thank Gregory J. Lawrence for assistance with sample preparation and helpful discussions. The work conducted by the U.S. Department of Energy Joint Genome Institute (https://ror.org/04xm1d337), a DOE Office of Science User Facility, is supported by the Office of Science of the U.S. Department of Energy operated under Contract No. DE-AC02-05CH11231.

---

**Supplementary Figure 1:**
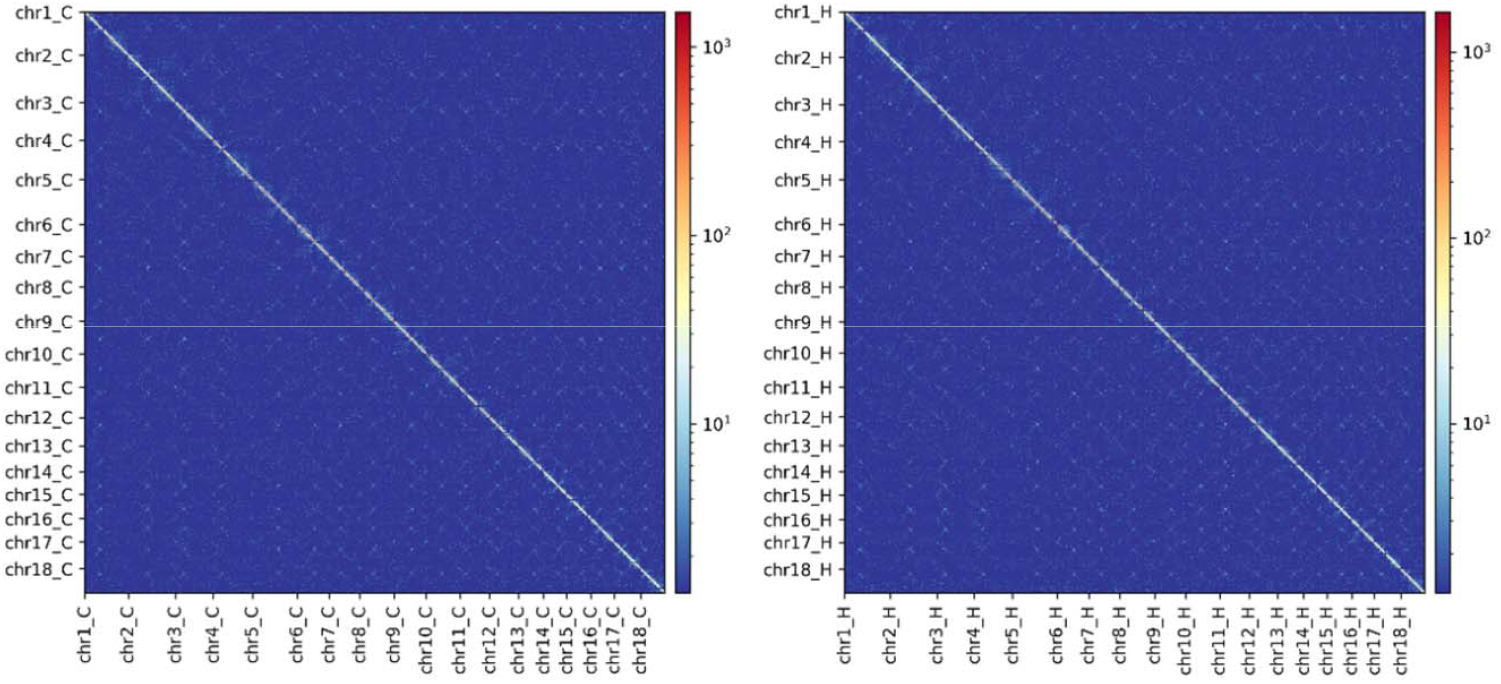
Hi-C contact maps of the two haplotypes show the location of the 18 centromeres as bowtie-like shapes.

**Supplementary Figure 2:**
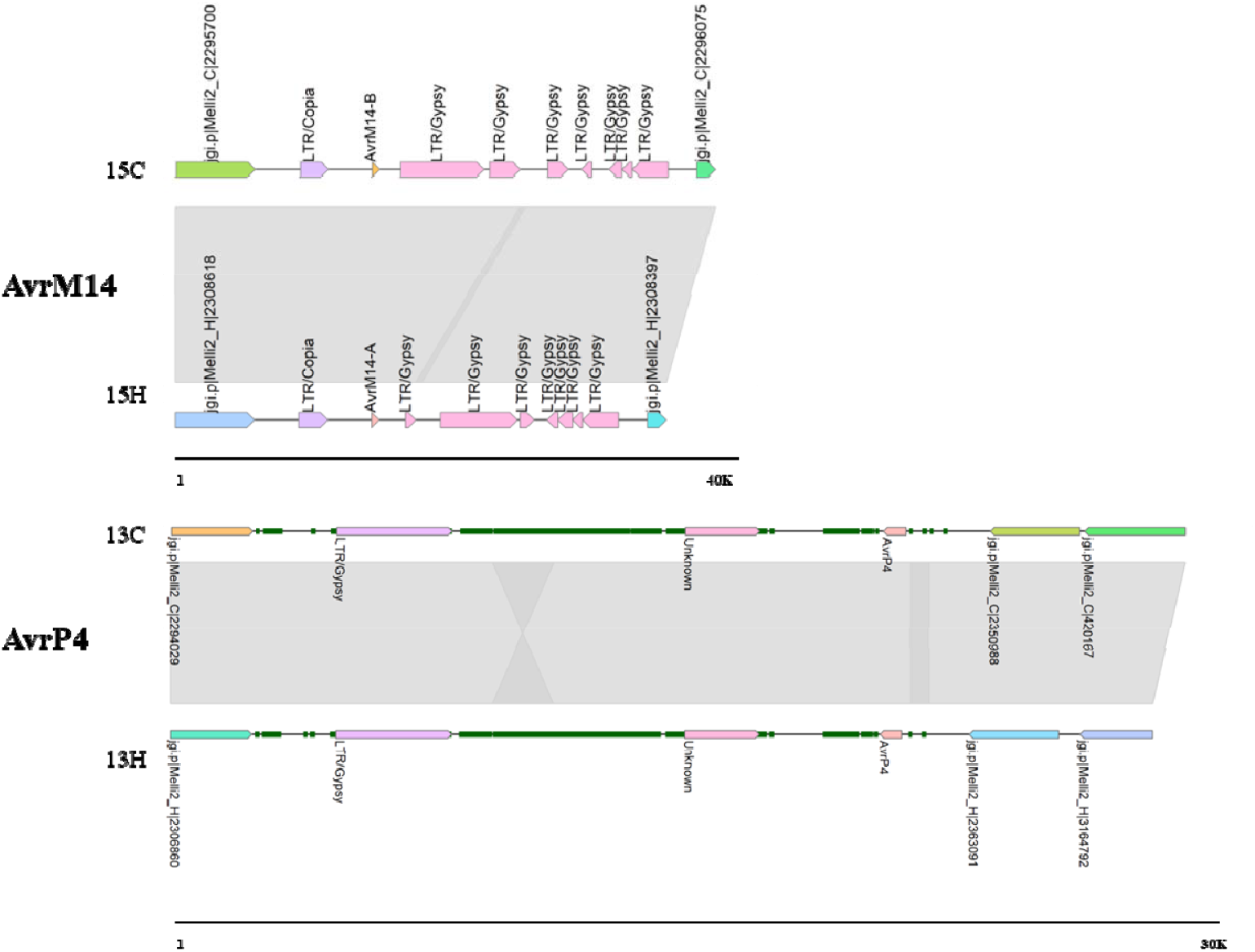
Conserved haplotype structures at the *AvrM14* and *AvrP4* loci in CH5. Figure shows synteny plots between the C and H haplotypes. Genes are labelled with their identifier. Repetitive elements are shown in green boxes. Separate scale bars are shown at the bottom of each locus. Shaded boxes connect haplotype regions of conserved sequence.

**Supplementary Figure 3:**
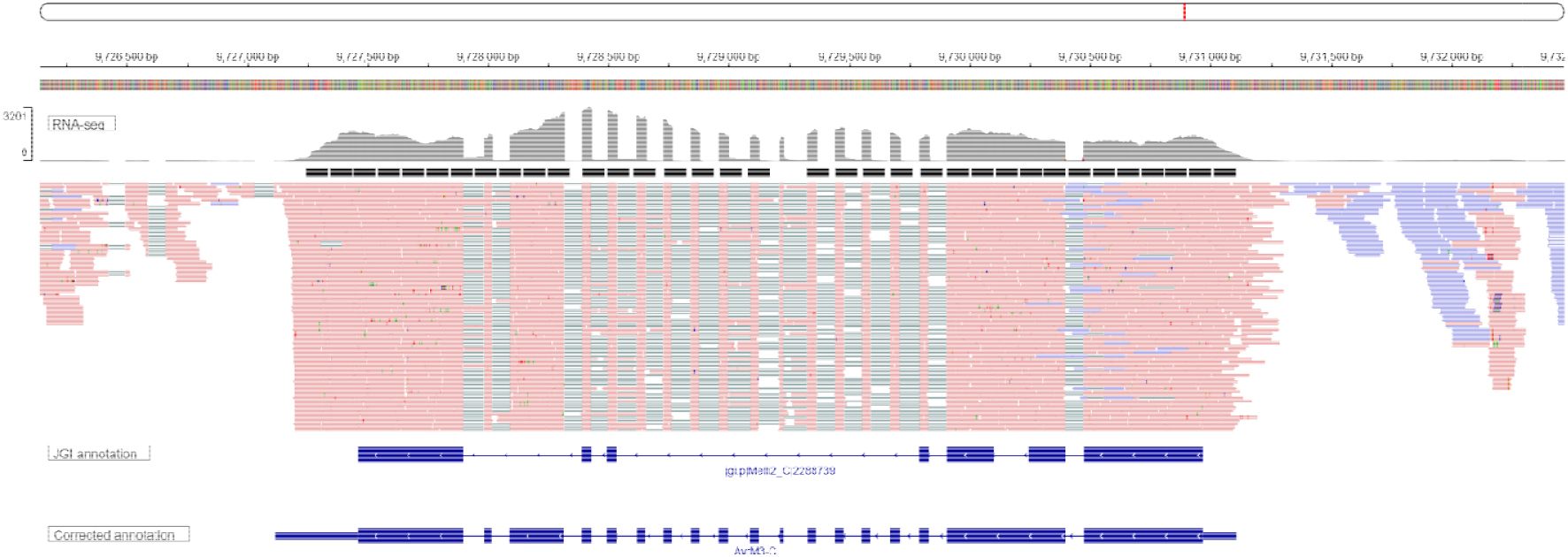
The AvrM3 candidate gene on the C haplotype was mis-annotated in the gene annotation. RNA-seq alignments are shown followed by the annotated gene and corrected gene model.

**Supplementary Figure 4:**
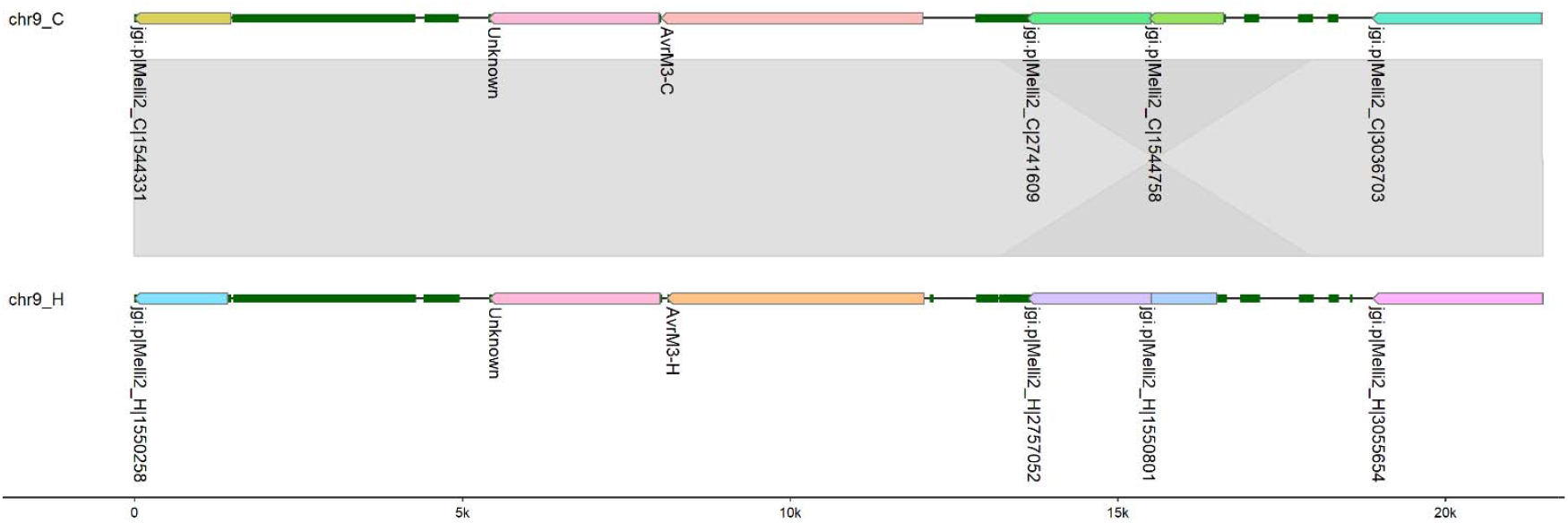
The AvrM3 locus is highly conserved between the two haplotypes. Figure shows synteny plots between the C and H haplotypes. Genes are labelled with their identifier. Repetitive elements are shown in green boxes. Separate scale bars are shown at the bottom of each locus. Shaded boxes connect haplotype regions of conserved sequence.

**Supplementary Figure 5:**
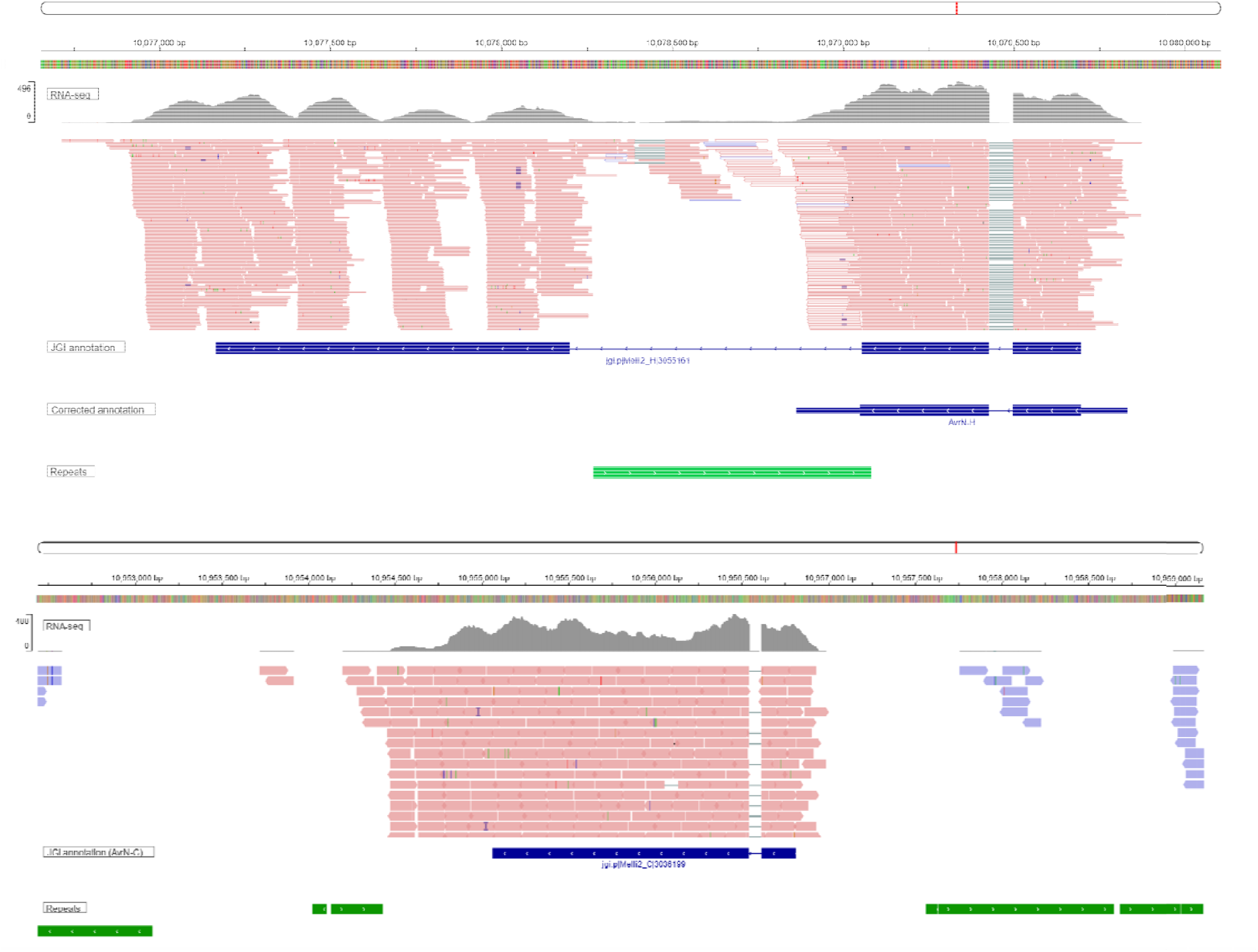
The AvrN allele in haplotype H was mis-annotated in the gene annotation. The AvrN-H locus is shown at the top and the AvrN-C locus is shown at the bottom. RNA-seq alignment do not support the intron in the annotated gene model for AvrN-H. The multi-mapped reads in the mis-annotated intronic region as well as the presence of an LTR Copia repeat element support the truncated H allele of *AvrN*. In comparison, the *AvrN*-C locus is correctly annotated and does not show a LTR Copia repeat insertion.

